# A Novel Class of Azoles with potent Anti-Leishmanial activity

**DOI:** 10.1101/2020.10.16.342329

**Authors:** Tarun Mathur, Manoj Kumar, Tarani Kanta Barman, V. Samuel Raj, Dilip J. Upadhyay, Ashwani Kumar Verma

## Abstract

**Objectives:** To investigate the anti-leishmanial activity of novel azole compounds against *Leishmania donovani*, which causes deadly visceral leishmaniasis disease.

**Methods:** A focused azole-based library was screened both against promastigotes and amastigotes forms of *L. donovani* strains in flat-bottomed 96-well tissue-culture plates and J774A.1 macrophages infected with *L. donovani*. The comprehensive screening azole-based library against *L. donovani* strains provided novel hits, which can serve as a good starting point to initiate hit to lead optimization campaign.

**Results:** Hits identified from azole-based library exhibited potent *in vitro* activity against promastigotes and amastigotes of *L. donovani*.

**Conclusions:** These potent novel azole hits could be a good starting point to carry out for further medicinal chemistry exploration for anti-leishmania program.

## Introduction

Leishmaniasis, is a vector-borne disease caused by obligate intracellular protozoa, and transmitted to human by the bite of a sand fly (1). Till date, 21 species of Leishmania have been identified and reported to cross the species barrier and cause infections in humans and cause three different clinical syndromes, visceral leishmaniasis (VL), cutaneous leishmaniasis (CL) and mucocutaneous leishmaniasis (MCL). Among these VL is most severe and remains a major public health problem worldwide (1). The human pathogenesis of *Leishmania* parasites is very complicated and dimorphic; the parasite can live and replicate in the gut of sandflies as flagellated forms (promastigote) or in mammalian cells (phagocytic, macrophages, fibroblast or dendritic cells) as aflagellated forms (amastigotes) (2).

According to the recent World Health Organization (WHO) report, leishmaniasis is the second-most deadly vector-borne parasitic disease, which represents a serious global health concern with a potentially fatal outcome (3). Worldwide, 12 to 15 million people are infected with *Leishmania* and 350 million are at risk of acquiring disease (4), with 1.5 to 2 million new cases every year (3). Among Leishmaniasis is endemic in over 98 countries and territories (5), but these majority of global VL cases are reported in India, Bangladesh, Nepal, and Brazil, whereas 60% of cases of CL occurred in South Asia and Middle east countries (4). Majority of VL cases are caused by *Leishmania donovani* and are life-threatening with a high fatality rate especially if left untreated (1). In addition, an overlap between the transmission areas of human immunodeficiency virus (HIV) and leishmaniasis has been reported in 35 countries (6), which has further complicated the clinical management of leishmaniasis (7).

The recommended current line of treatment for VL still includes old drugs such as, sodium antimony gluconate (SAG), Pentostame, Meglumine antimoniate (Glucantime), Amphotericin B and its lipid formulation AmBisome, Pentamidine and Miltefosine (8, 9). Unfortunately, current line of treatment are associated with multiple drawbacks; high toxicities, poor patient compliance (long-term parenteral administration) etc. In addition, high cost of AmBisome and Miltefosine treatment and emergences of resistant strains to current line of treatment has further limit their use in clinics (10–12). Though, some vaccine candidates are identified along the years, but till date, none are available for clinical use in human to save the lives of infected persons (13). Given the limitations of current anti-leishmanial drugs, including the availability of efficacious drugs, cost of treatment (drugs and hospitalization), toxicities, and growing resistance strains, has created an unmet medical need for affordable and novel drugs candidate against VL, in the endemic area to promote better healing and save lives of infected patients (10, 14). Considering the success of oral antifungal azole drugs such as Ketoconazole and Itraconazole and their reported potent activities against *Leishmania* sp., (15–18), identification of such novel azole compounds would be a significant advance in *Leishmania* treatment that would further aid in patient’s compliance and disease control/elimination efforts. Azoles have been known to inhibit the cytochrome P 450-mediated 14α-demethylation of lanosterol, blocking ergosterol synthesis and causing accumulation of 14α-methyl sterols, which proves lethal to the organism (19, 20). In view of the great success of azole antifungal drugs in last three decades, we planned to assess potential of in-house azoles library against both, promastigote and amastigotes forms of *Leishmania*. These novel compounds were synthesized for antifungal discovery program and many of them demonstrated activity superior than known azole-anti fungal drugs against various fungal isolates (21, 22). A hit identification campaign using Ranbaxy’s azole library was planned with the objective to identify a good starting point for initiating a hit-to-lead medicinal chemistry efforts for anti-leishmania program.

## Material and methods

### *L. donovani* strains and compounds

Clinical *Leishmania* isolates were obtained from All India Institute of Medical Science (AIIMS), New Delhi, and standard strains were purchased from the American Type Culture Collection (ATCC, Manassas, VA, USA). *L. donovani* AG 83 was obtained from ATCC and *L. donovani* 2001-S, and *L. donovani* were received from AIIMS. Promastigotes were grown in Hepes-buffered medium M-199 (Gibco Rockville, MD) supplemented with 10% heat inactivated fetal bovine serum at 22°C. Amastigotes were collected from spleen of an infected hamster and maintained in Schneider’s Drosophila medium (SM medium, Gibco Rockville, MD) at 37°C/ 5% CO_2_. J774.1 macrophages were maintained in RPMI 1640 medium (Hyclone, USA) supplemented with 10% fetal bovine serum at 37°C in 5% CO_2_ in a humidified atmosphere. The reference antileishmanial agents [Amphotericin B (Sigma), SAG (Pentostam), Miltefosine (Asta indica) and azoles (Miconazole, Ketoconazole, Fluconazole and Itraconazole) were obtained from commercial sources. 500 RBx azole library compounds were obtained from Ranbaxy’s compound library and stock solutions were prepared in dimethyl sulfoxide (DMSO) at 10 mg/ml. For the *in vitro* screening, the compound was further diluted using appropriate medium.

### Drug susceptibility assay against promastigotes

For the screening of library, an early log phase promastigote (2 × 10^6^ promastigotes per ml) were seeded in 96-well flat-bottom plastic tissue-culture plates (Nunc Int., Denmark) and incubated at 22^°^C for 72 hours in presence or absence of compounds. After the incubation, the viability of promastigotes was checked using the Alamar blue as described earlier (23–25). The concentrations inhibiting parasite growth by 50% (GI_50_) was calculated from the dose response curve by using Graph Pad, Prism (San Diego).

### Drug susceptibility assay against amastigotes

Drug susceptibility of amastigotes was performed using the J774A.1 monocyte-macrophage mouse line. Briefly, J774A.1 macrophages were seeded at 4×10^5^ macrophages/well in RPMI with 10% FBS in sterile 16-chamber slides (Nunc Int., Denmark). After incubation, 100 μl of suspension containing amastigotes (4.2 × 10^6^/ml) were added to each well, at a ratio of 10 parasites per macrophage. After 24 hours, the free amastigotes were removed and intra-macrophagic amastigotes were treated with various concentrations of the test compounds for 48 hours of incubation. Following incubation, the cells were fixed with methanol and stained with 10% Giemsa (Hi Media, India). Infected and non-infected macrophages were counted in the control cultures and those exposed to the serial drug dilution. Each experiment was performed in triplicate. The results were expressed as percent reduction in parasite burden compared to non-infected control wells, and the GI_50_ was calculated by linear regression analysis using Graph Pad prism.

## Results and discussion

Sterol biosynthesis in fungi and *Leishmania* resembles in synthesizing 24-substituted sterols such as ergosterol (ergosta-5, 7, 24(28)-trien-3β-ol (5-dehydroepisterol) and sterol biosynthesis appear to proceed via lanosterol (26). Azoles are known to impair ergosterol production in fungi by inhibiting the cytochrome P 450-dependant 14α-demethylation of lanosterol or 24-methylene-dihydro lanosterol (27). Considering the potent anti-Leishmania activity of azoles, Ketoconazole, Itraconazole and Fluconazole have undergone several CL and VL trials with equivocal results (8). Itraconazole and other azoles have demonstrated anti-leishmanial activity, especially in cutaneous leishmaniasis and some reports of their usage in VL (21, 28, 29). To further validate these reports, we screened azole based known antifungal drugs such as Itraconozole, Fluconazole, Miconazole and Ketoconazole and they exhibited 50% inhibitory concentrations (GI_50s_) in the range of 16-64 μg/ml against promastigotes of *L. donovani* (14, 16). Table-1. After screening the in-house azole-based compounds library, two compounds (RBx-0831 and RBx-0091) were identified as an inhibitor of *Leishmania* promastigotes (GI_50_ 4-20 μg/ml. The antileishmanial drug, SAG did not show any activity against promastigotes (GI_50_ >64 μg/ml) whereas, promastigotes were found to be highly susceptible to Amphotericin B (0.2 μg/ml) and Miltefosine (1.5 to 7 μg/ml). These results have been summarized in Table-1.

**Table 1.** *In vitro* activity of compounds against *L. donovani*

| Test Compounds | Susceptibility [GI<sub>50</sub> (μg/ml)]<sup>a</sup> |  |  |  |  |  |  |
| --- | --- | --- | --- | --- | --- | --- | --- |
|  | Promastigotes |  |  | Amastigotes |  |  | Cytotoxicity |
|  | *L. donovani* Clinical strain. | *L. donovani* AG 83 | *L. donovani* 2001-S | *L. donovani* Clinical strain | *L. donovani* AG 83 | *L. donovani* 2001-S | J774.1 macrophage |
| RBx-0831 | 6 ± 0.02 | 4 ± 0.02 | 5.6 ± 0.03 | 4.0 ± 0.05 | 2.5 ± 0.02 | 1.4 ± 0.1 | >50 |
| RBx-0091 | 16 ± 0.20 | 16 ± 0.10 | 20 ± 0.25 | 5.6 ± 0.2 | 5 ± 0.05 | 6.1 ± 0.2 | >50 |
| Fluconazole | >64 ± 0 | >64 ± 0 | >64 ± 0 | >32 ± 0 | >32 ± 0 | >32 ± 0 | >50 |
| Itraconazole | >64 ± 0 | >64 ± 0 | >64 ± 0 | >32 ± 0 | >32 ± 0 | >32 ± 0 | >50 |
| Miconazole | 32 ± 1.10 | 32 ± 1.25 | 64 ± 1.35 | >32 ± 0 | 32 ± 1.25 | 16 ± 1.2 | >50 |
| Ketoconazole | 16 ± 0.45 | 16 ± 0.20 | 17 ± 0.4 | 16 ± 1.2 | 28 ± 1.5 | >32 ± 0 | >50 |
| SAG | >64 ± 0.5 | >64 ± 0.4 | >64 ± 0.5 | 4.5 ± 0.2 | 6.0 ± 0.3 | 5.1 ± 0.6 | >50 |
| Amphotericin B | 0.02 ± 0.01 | 0.02 ± 0.02 | 0.02 ± 0.01 | 1.5 ± 0.2 | 0.06 ± 0.02 | 0.2 ± 0.2 | >50 |
| Miltefosine | 7.0 ± 0.01 | 1.4 ± 0.02 | 4.5 ± 0.01 | 1.99 ± 0.18 | 0.5 ± 0.02 | 2.06 ± 0.14 | >50 |

<sup>a</sup> Mean GI<sub>50s</sub> ± standard deviations of the results from three separate assays. SAG, sodium antimony gluconate.

RBx-0831 and RBx-0091 exhibited GI_50_ of 2.5 and 5 μg/ml, respectively against intracellular amastigotes whereas Fluconazole and Itraconazole did not exhibit any activity against *L. donovani* amastigotes up to 64 μg/ml. SAG, Amphotericin B and Miltefosine exhibited GI_50_ of 6.0, 0.06 and 0.5 μg/ml, respectively. The *in vitro* potential of RBx-0091 was found similar to Miltefosine and better than SAG and Ketoconazole against amastigotes, while RBx-0091 exhibited better *in vitro* potential against both promastigotes and amastigotes form of parasites than Fluconazole, Itraconazole, Miconazole, and Ketoconazole. As expected, Amphotericin B and Miltefosine showed potent activities, both promastigotes and amastigotes forms of parasites in our screening; in contrast, fluconazole, itraconazole, miconazole, and ketoconazole exhibited poor anti-Leishmania activities. The screening results of the above standard drugs were found in agreement with earlier reports (30).

The clinical impact of VL is very severe and response to recommended therapeutic approaches is poor (11, 21). Although, the second-line of treatment using Amphotericin B is quite efficacious for treating these antimony-resistant patients but its usage is limited due to serious side effects, resistance and poor patient compliance (10). Although, alternative treatment for VL is currently available but these remain largely inaccessible to poor patients because of their high cost. The discovery of potent novel azoles-based hits against *Leishmania* has opened up a new opportunity for further medicinal chemistry exploration which may provide a cost-effective, safe and oral treatment against this disease. Further evaluation of these hits for pharmacokinetics studies and in the hamster model for *in vivo* efficacy, which closely mimics the progression of human VL, is warranted. Presently, we are in the process of evaluating results obtained from our study in terms of understanding the mechanism of action, profiling against different resistant isolates, and medicinal chemistry exploration to further optimize these novel hits against *L. donovani*.

**Figure 1.**
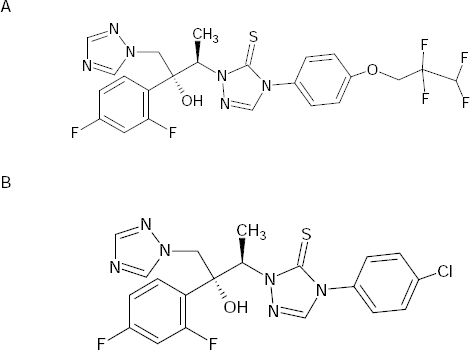
Structure of RBx 0831 (A) RBx 0091 (B).

## Acknowledgements

This work was made possible by a fund from Ranbaxy research Laboratories. The findings herein reflect the work and are solely the responsibility of the authors. We thank Dr. Pradip Kumar Bhatnagar for his critical review of this manuscript.

## Transparency declarations

‘None to declare’

